# Thermodynamics and kinetics of aggregation of flexible peripheral membrane proteins

**DOI:** 10.1101/2021.04.09.439228

**Authors:** Mohsen Sadeghi, Frank Noé

**Affiliations:** Department of Mathematics and Computer Science, Freie Universität Berlin, Arnimallee 12, 14195 Berlin, Germany

## Abstract

Biomembrane remodeling is essential for cellular trafficking, with membrane-binding peripheral proteins playing a key role in it. Significant membrane remodeling as in endo- and exocytosis is often due to aggregates of many proteins with direct or membrane-mediated interactions. Understanding this process via computer simulations is extremely challenging: protein-membrane systems involve time- and lengthscales that make atomistic simulations impractical, while most coarse-grained models fall short in resolving dynamics and physical effects of protein and membrane flexibility. Here, we develop a coarse-grained model of the bilayer membrane bestrewed with rotationally-symmetric flexible proteins, parametrized to reflect local curvatures and lateral dynamics of proteins. We investigate the kinetics, equilibrium distributions, and the free energy landscape governing the formation and break-up of protein clusters on the surface of the membrane. We demonstrate how the flexibility of the proteins as well as their surface concentration play deciding roles in highly selective macroscopic aggregation behavior.

---

Membrane-bound peripheral proteins are essential to a wide range of biological functions, especially when they are capable of sensing/inducing membrane curvature. These proteins are often found in regions of the plasma membrane that undergo significant conformational changes in processes such as tubulation, exo/endocytosis, and fission/fusion. The same also holds for pathogens such as Shiga and Cholera toxins [1–3]. Membrane remodeling leading to internalization of these toxin can progress without relying on active cell machinery [2, 4], highlighting the importance of understanding the protein-membrane mechanical interplay. Considering the rather large bending stiffness of bilayer membranes (∼20 kT), macroscopic membrane remodeling involves significant energy expenditure that is beyond the action of any single membrane-binding protein. Thus, independent of binding and curvature-generation mechanism [5–8], membrane remodeling necessitates cooperative action of an assembly of peripheral proteins [9, 10].

Due to limitations of simultaneously resolving time and space at the nanoscale with experimental appraoches, physics-based computer models are a key technology to gain insights into structural and spatiotemporal mechanisms. However, physics-based models of the peripheral protein clusters are challenged by multiscale nature of the problem: on one hand, a simulation model must provide enough detail to resolve the relevant molecular effects, on the other hand, it needs to cover time- and lengthscales not simultaneously achievable by atomistic models.

For example, lipid reorganization has been indicated when these proteins are in close proximity [11]. Yet, macroscopic dynamics of protein clusters implies the existence of long-range membrane-mediated interactions (see [12] and references within), whose nature has been the subject of numerous biophysical studies [13–19]. With experimental data lacking, simulation tools have proved essential in the study of these interactions and the resulting organization and phase separation [20– 25]. Non-symmetric proteins such as crescent-shaped BAR-domains induce strong membrane-mediated interactions resulting in salient aggregation and structuring on the membrane [10, 26–28]. However, the interactions and cooperativity of rotationally symmetric proteins, such as STxB, seem to rely on the much more subtle fluctuation-induced forces [12–16, 25]. These so-called thermal Casimir interactions bring proteins that suppress membrane fluctuations together to minimize their adverse entropic effect. Despite several studies, and a multitude of hypotheses, reliable quantitative results that offer insight into the cooperative action of these proteins leading to macroscopic aggregation are scarce. Also, whilst the kinetics of membrane and peripheral proteins are of utmost importance in remodeling processes [29], developing a model that incorporates lateral kinetics of the proteins on the same grounds as the large-scale kinetics of the membrane and solvent is still challenging.

Here we develop a coarse-grained model of flexible membrane-bound proteins, diffusing on and interacting with a dynamic membrane. The model is fully particle-based so as to facilitate the integration with particle-based simulations of cellular kinetics [30, 31], and particularly with interacting-particle reaction dynamics (iPRD) [32–35]. We study the dynamics of protein aggregation and uncover slow kinetics in large-scale simulations. We develop a theoretical model describing this process, and yielding information on cluster formation/break-up kinetics as well as the chemical potential governing particles moving between clusters. Using histogram reweighting, we also obtain the free energy landscape of protein aggregation. We suggest a pathway for spontaneous aggregation similar to the nucleation curve, and discuss the opposing effects of protein stiffness.

## MODELING

### Particle-based model of flexible membranes and membrane-bound proteins

We have developed an extended version of the particle-based membrane model of ref. [39] (Supplementary Information). We have chosen a lattice parameter of 6.5nm, comparable with the radius of the STxB protein, to represent a membrane that allows tightly packed clusters of proteins. Membrane particles can be tagged to represent where peripheral proteins are bound (Fig. 1a). All effects corresponding to the presence of a peripheral protein are incorporated in a locally masked force field.

**Figure 1:**
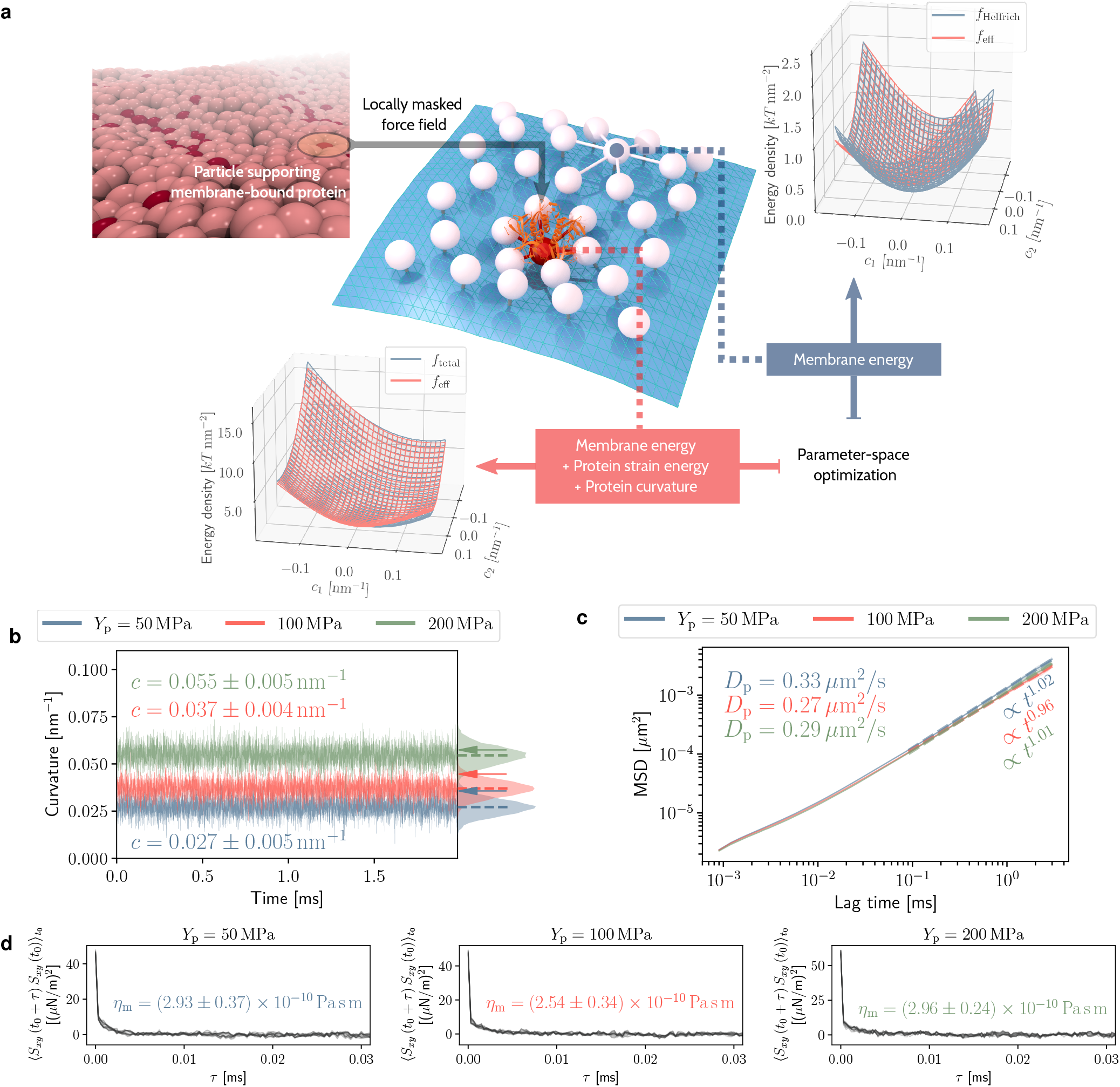
Dynamic model of membrane and peripheral proteins: (**a**) Mesoscopic membrane model with selected particles “tagged” as supporting peripheral membrane-bound proteins. Where a protein is bound to the membrane, a locally modified force field is applied (masking of the force field). Force field parameters are optimized to reproduce the Helfrich energy density for regular particles, plus an additional strain energy corresponding to the deformation of peripheral proteins for tagged particles (Supplementary Information). The two plots compare, over the given range of membrane curvatures, the energy densities of the theoretical models with the effective energy densities resulting from particle-particle interactions, when the optimized force field is used. In the lower plot, results are given where the Young modulus of *Y*_p_ = 100MPa, and an intrinsic curvature of *c*_0_ = 0.08nm^−1^ are attributed to the peripheral proteins. (**b**) Local membrane curvature in the vicinity of peripheral proteins with the given stiffness and the intrinsic curvature of *c*_0_ = 0.08nm^−1^, obtained from simulations of∼ 325×325 *μ*m^2^ membrane patches containing 10 % surface concentration of bound proteins. Distribution of instantaneous curvature values is shown on the right. Dashed lines pinpoint the mean of each distribution, while the shaded regions around the time series correspond to one standard deviation. The arrows point to the estimated curvatures from the parameter-space optimization, obtained from energy distributions in curvature space shown in (a). (**c**) Mean-squared displacement for lateral motion of proteins. Dashed lines represent power-law fits to the shown range of lag times. Values of the corresponding exponents of time *t* and the effective long-time diffusion coefficients are color-coded for the given protein stiffnesses. (**d**) Time-correlation functions of the in-plane shear stress of the membrane, based on which, membrane surface viscosity (*η*_m_) is obtained from the corresponding Green-Kubo relation [36–38].

### Effective membrane curvatures

Distribution of effective energy density (*f*_eff_) in membrane + protein regions (Fig. 1a inset plots) yields an initial estimate of the local membrane curvatures, depending on the intrinsic curvature of proteins as well as their stiffness (Supplementary Information). The actual curvature observed in simulations deviates because: (i) the parametrization scheme is only applied considering the nearest neighborhood of each particle, and (ii) the local masking of the force field is achieved through Monte Carlo moves to preserve detailed balance (Supplementary Information). Not surprisingly, stiffer proteins lead to more pronounced curvatures (Fig. 1b) with the energy-based estimate providing an upper bound (Fig. 1b, arrows). The wide range of curvatures facilitates modeling a variety of proteins. Previous molecular dynamics simulation of STxB bound to Gb3 lipids predicted inward membrane curvatures of 0.034 ± 0.004nm^−1^ for saturated and 0.035 ± 0.003nm^−1^ for unsaturated acyl chains on the lipids [40]. Also, using curvature sorting of Cholera toxin subunit B (CTxB), the induced membrane curvature has been experimentally determined to be 0.055 ± 0.012 nm^−1^ [41].

### Kinetics of membrane and proteins

In order for the predicted timescales and dynamical mechanisms to be reliable, the model must faithfully capture the kinetics of the membrane and membrane-bound proteins. With the current model, including hydrodynamic coupling, we have previously established that for the kinetics of out-of-plane membrane fluctuations [36]. Here we investigate the lateral diffusion of protein-bound particles (Fig. 1c) and subsequently calibrate it to the experimental range of values via tuning the frequency of bond-flipping moves and the in-plane mobility of particles (Supplementary Information and [36, 39]).

The mean-squared displacement (MSD) of proteins follows the familiar three-step regime [42]: fast diffusion for short time intervals, followed by subdiffusive behavior, and finally, the asymptotic normal diffusion for large lag times, with MSD proportional to 4*D*_p_*t* (Fig. 1c). Consulting MSD values, switching to normal diffusion happens as soon as particles leave their immediate neighborhood. Protein diffusion coefficients compare well with the experimental values. Using fluorescence recovery after photobleaching (FRAP) assays, Tian and Baumgart measured a diffusion coefficient of 0.35 ± 0.09 *μ*m^2^ s^−1^ for CTxB proteins [41]. Also, Day and Kenworthy similarly measured the diffusion coefficient of STxB proteins as ∼0.5 *μ*m^2^ s^−1^[43]. It is noteworthy that multivalent glycolipid-binding proteins such as CTxB or STxB diffuse extremely slowly compared to their lipid-anchored counterparts [44].

Irrespective of the protein stiffness, time-correlation functions of membrane’s in-plane shear stress decay very rapidly and do not show any long-term memory effects (Fig. 1d). Surface viscosity, especially considering the error margins, is almost independent of protein stiffness (Fig. 1d). The experimental value for the surface viscosity of the membrane is in the 10^−10^ – 10^−7^ Pa sm range [45–48]. While molecular dynamics simulations with coarse-grained force fields usually result in much lower viscosities of 10^−13^ – 10^−11^ Pa sm [49, 50]. Despite being highly coarse-grained, thanks to the anisotropic stochastic dynamics (Supplementary Information), we could successfully produce values in much better agreement with experiments.

## RESULTS

### Dynamics of protein aggregation

Formation of stable protein clusters relies on cooperative action, and possibly multi-body interactions, making it a rich and complicated process [9, 25]. Among the significant body of work on thermodynamics and statistical mechanics of particle aggregation (see for example [51] and the references within), we have chosen the kinetic theory based on the following reaction [52–54],

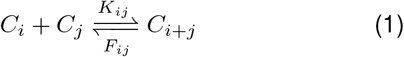

where *C*_*k*_ denotes a cluster of *k* particles, and *K*_*ij*_ and *F*_*ij*_ are the corresponding coagulation (formation) and fragmentation (break-up) rates. This model leads to an equilibrium cluster size distribution, while allowing for the formation of one macroscopic cluster (Methods section “Cluster size distribution”).

We analyze 3 ms of simulation data, presenting an abundance of these formation/break-up events (Fig. 2a). The cluster growth follows a rather slow dynamics, especially for stiffer proteins (Fig. 2b). The measured timescale, *τ*, can be compared to the diffusion times measured for STxB bound to giant unilamellar vesicles (GUV’s) [24]. With the diffusion coefficients measured here, for *Y*_p_ = 200MPa proteins, proteins can travel distances of ∼35nm in between coagulation/fragmentation events. This is comparable to the diameter of a cluster formed by 30 proteins (compare to Fig. 2c).

**Figure 2:**
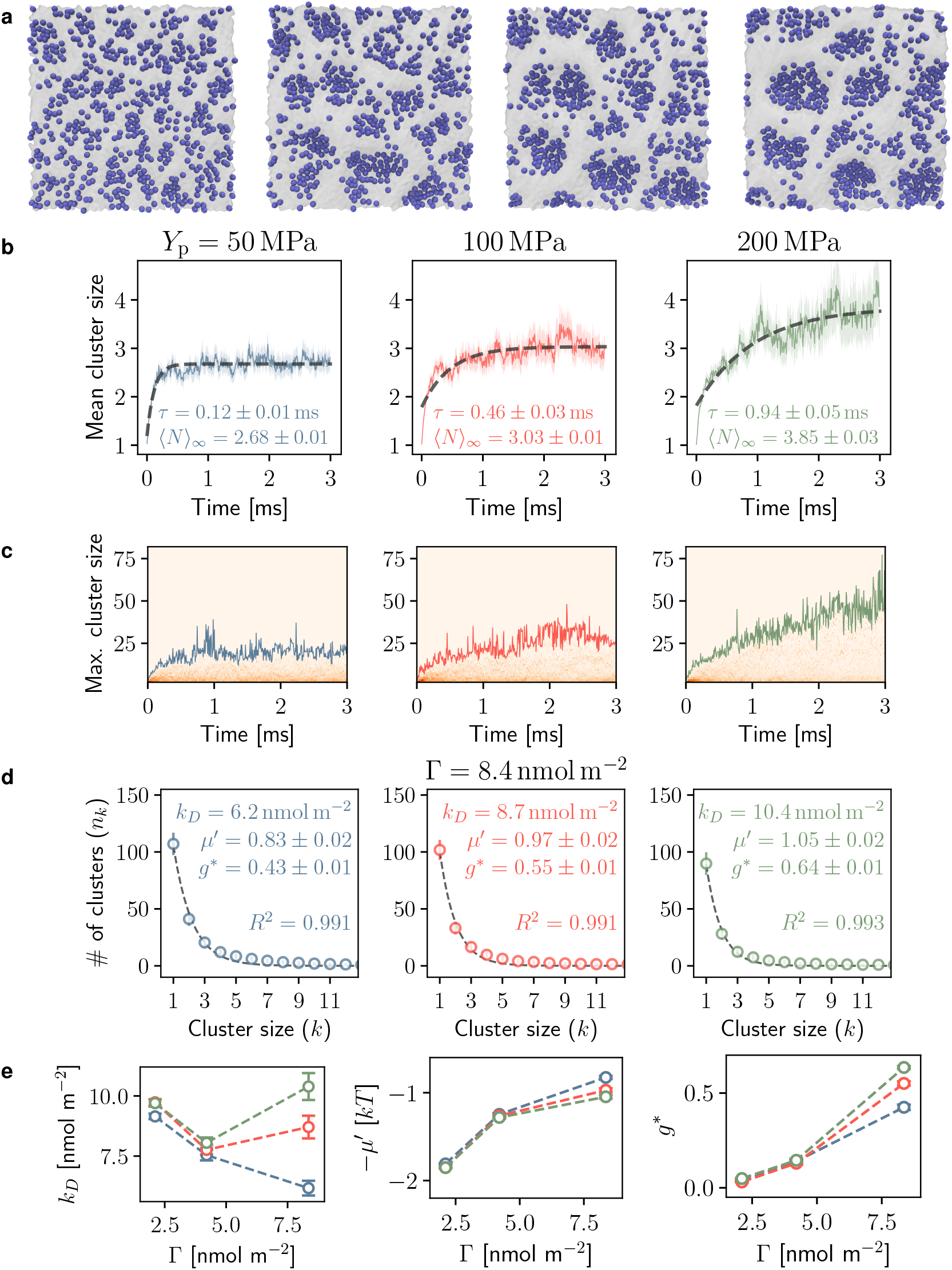
Dynamics of protein clusters on the surface of the membrane: (**a**) Simulation snapshots showing the formation and break-up of peripheral protein clusters for the high protein surface concentration of Γ = 8.4nmolm^−2^ (Supplementary Movie) (**b**) Time evolution of the weighted mean cluster size (number of particles in a cluster) for peripheral proteins with the given stiffnesses, with the same concentration as in (a). Dashed lines are fits of an exponential saturation model ⟨ *N*⟩_*t*_ = ⟨*N*⟩_∞_ + (⟨ *N*⟩_0_− ⟨ *N* ⟩_∞_)exp (−*t/τ*), with values of timescale *τ*, and equilibrium mean cluster size ⟨ *N* ⟩_∞_, given in each case. (**c**) Maximum cluster size achieved during the simulation. The background of each plot shows the evolution of the histogram of cluster sizes. (**d**) Equilibrium distribution of cluster size for the given protein surface concentration. Dashed lines are fits of Eq. (6). Best fit parameters and the *R*^2^ score of the fit are given on the plots. (**e**) From left to right: dissociation constant, chemical potential, and size of the macroscopic cluster, obtained from fits of the most probable cluster size distribution, as a function of the surface concentration of proteins. Colors correspond to protein stiffness, as in (a).

The most probable cluster size distribution predicted by the kinetic theory (Eq. (6)) very well fits our simulation results (Fig. 2d). Comparing Eq. (6) with the probability distribution associated with the grand canonical ensemble, *p* ∝ exp (*μN/kT*), suggests −*μ*′ to serve as the chemical potential (in units of *kT*) for exchange of particles between clusters. Lower chemical potential hints at more stable aggregates. We observe that at low protein surface concentrations (Γ) protein stiffness only slightly affects the chemical potential. But at higher Γ, for which the snapshots in Fig. 2a clearly show the formation of large clusters, chemical potential depends on protein stiffness, with stiffer proteins tending toward more stable clusters (Fig. 2e). The maximum cluster size is also achieved for the stiffest proteins (Fig. 2c). Yet, comparing the numerical values of *μ*′ points at the interesting fact that from the free energy point of view, addition of particles to clusters at lower concentrations is more favorable.

We found the dissociation constant (*k*_*D*_) to change with Γ in a non-trivial manner. At low Γ, *k*_*D*_ decreases with concentration for all proteins, i.e. the reaction (1) shifts towards more clusters forming than fragmenting (Fig. 2e). But at high Γ, protein stiffness exerts a significant influence. While so far we have observed *Y*_p_ = 200MPa proteins to be successful in forming large clusters at high Γ, they have the highest *k*_*D*_, pointing to more fragmentation events occurring. It is noteworthy that the model has a singular *k*_*D*_ parameter, independent of cluster sizes *i* and *j* in Eq. (1). Thus, these fragmentation events most probably correspond to small unstable clusters.

The size of the macroscopic cluster predicted by the theoretical model is highlighted by the *g*\* parameter in Eq. (6). *g*\* generally grows with Γ, increasing drastically to 43 – 64 % for Γ = 8.4 nmolm^−2^. As expected, this increase is most noticeable for *Y*_p_ = 200MPa, hinting at a phase transition at high surface concentration.

### Free energy landscape of protein aggregation

Free energy landscape of the aggregation process offers the best outlook on the stability of protein clusters and the nature of phase transitions. To this end, we have performed equilibrium simulations with different surface concentration of peripheral proteins. We use a reaction coordinate *q*, corresponding to the continuous dispersion ↔ aggregation transition, and use histogram reweighting for free energy estimation (Methods section “Free energy calculation”). Clear separation of the histogram of *q* values for simulation frames where the mean cluster size varies minutely implies this reaction coordinate to be suitable for discriminating dispersed and aggregated states (Fig. 3b).

**Figure 3:**
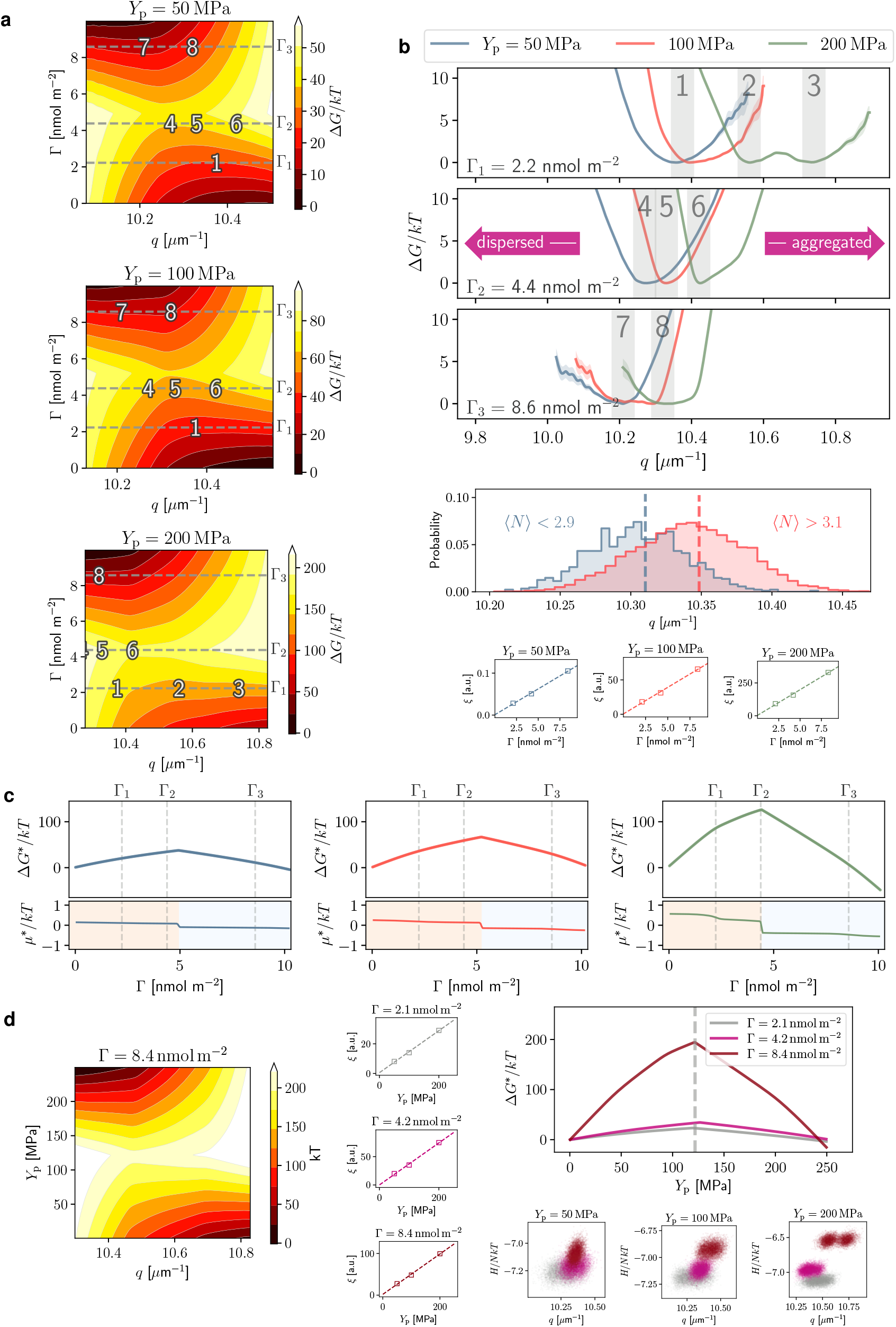
Free energy of protein aggregation: (**a**) Free energy landscape along the aggregation reaction coordinate, *q* (Eq. (7)) and surface concentration, Γ. (**b**) Top: Free energies at surface concentrations Γ_1_ and Γ_2_, marked by gray dashed lines in (a). Shaded regions coincide with numbered states in (a). Middle: the distribution of the reaction coordinate *q*, where the mean cluster size matches the given criteria. Bottom: correlation between the inferred bias, *ξ*, and Γ (Methods section “Free energy calculation”). (**c**) The marginalized free energies (top plots) and chemical potentials (bottom plots) versus surface concentration, Γ. Colors correspond to the protein stiffness similar to (b) (**d**) Left: free energy landscape along *q* and protein stiffness, *Y*_p_. Middle: correlation between the inferred bias, *ξ*, and protein stiffness. Right top: marginalized free energy as a smooth function of protein stiffness for different surface concentrations. Right bottom: joint distribution of *q* and mean energy for the given protein stiffness and different surface concentrations (colors correspond to the legend of the middle plot).

The free energy landscape strongly depends on the stiffness and surface concentration of proteins (Fig. 3). To avoid confusion, we need to distinguish between the aggregation described by *q*, and the global surface concentration, Γ. While high Γ indicates a large copy numbers of proteins being present in the system, it does not imply the formation of tight clusters. This clarifies the seemingly contradictory observation that increasing Γ results in the free energy minima generally shifting toward more dispersed states (Fig. 3a). This shift is more drastic for the *Y*_p_ = 50MPa and 100MPa proteins, compared to *Y*_p_ = 200MPa (Figs. 3a and 3b, numbered states can be used as reference between plots). For the stiffest proteins, coexistence between aggregated phases “2” and “3” is favorable at low Γ, with the semi-aggregated state “6” existing in the mid-range Γ.

Generally, the free energy landscape over *q* and Γ has a saddle point, separating the basins corresponding to low and high concentration states. Marginalizing the free energy along *q* reveals a behavior very well resembling the classical nucleation curve (Fig. 3c). There exists a critical surface concentration beyond which addition of proteins results in free energy reduction (Fig. 3c). This is best demonstrated by the chemical potential *μ*\* = *∂G*\**/∂N* (Fig. 3c). The switch to negative chemical potential denotes the onset of spontaneous cluster growth. This switch happens at increasingly lower surface concentration as the stiffness increases. However, the value of *μ*\* compared to *kT* is only significant for *Y*_p_ = 200MPa proteins (Fig. 3c).

At high surface concentration, free energy landscape along *q* and protein stiffness uncovers how increasing the protein stiffness is favorable beyond *Y*_p_∼100MPa. This distinction is less apparent at lower concentrations (Fig. 3d). The joint distribution of *q* and internal energy shows how a higher energy but more aggregated state emerges as the protein stiffness increases (Fig. 3d).

## DISCUSSION

We have presented a dynamical model of biomembranes with flexible membrane-bound proteins to study their aggregation behavior. The model exhibits membrane surface viscosity and protein diffusion coefficients in close proximity of experimental values, while allowing for a range of membrane curvatures and protein flexibilities. For example, for the stiffest proteins that we considered, the local membrane curvature in our model is 0.055nm^−1^, closely matching the data obtained from curvature sorting of CTxB proteins [41]. This model is suitable for large-scale simulations elucidating collaborative actions and macroscopic phenomena.

The model is purposefully void of a direct interaction between proteins. Nevertheless, peripheral proteins aggregate on the membrane via interactions that are evi-dently of membrane-mediated nature (Fig. 2a and Supplementary Movie).

We have obtained two estimates of the chemical potential. The first corresponds to existing proteins joining clusters (∼ 1 *kT/*particle), while the second describes the free energy change due to new proteins entering the system (∼ 0.5 *kT/*particle). Both these findings point to the fact that the magnitude of attractive membrane-mediated interactions are small [9]. Yet, the collaborative effect of these interactions seem to suffice to form clusters, albeit in a dynamical and sometimes loosely-bound fashion (Fig. 2a). Based on free energy estimates, we found that favorable cluster growth at higher Γ necessitates smaller values of *q*, i.e. the most stable configurations at high surface concentration might not be tightly bound macroscopic clusters, but either a dynamic state of loose aggregates or clusters that repel and reorganize in a relaxed lattice when they reach a certain size. Evident for both behavior can be observed in our simulations (Fig. 2a and Supplementary Movie). The rather high nucleation barrier for *Y*_p_ = 200MPa proteins also hints at large energetic penalty, possibly due to repulsive interactions, before high-concentration coalesced states become possible.

Higher protein stiffness, which in our model also includes stronger membrane-binding, results in larger fluctuation-based attraction, but also enhances repulsions originating from membrane bending energy. This contrariety explains the rich aggregation behavior observed. Beyond a critical surface concentration, spontaneous aggregation occurs (Figs. 3c and 2a). This dependence on surface concentration has been previously observed in experiments on GUV’s [24]. We also observed that for stiffer proteins, the dissociation constant increases at high surface concentration (Fig. 2e). We believe this points to small cluster being dispersed at a higher rate when the surface concentration is high.

Formation of large protein clusters results in macroscopic membrane conformation, the onset of which can be observed in Supplementary Movie. The sequence of events in membrane remodeling pathway and the accompanying free energy landscape are well within reach with the current model. The approach laid out here paves the way for direct comparison of simulation and experiment to study processes such as clathrin-independent endocytosis of toxins or signaling molecules. It also offers a quantitative tool for designing finely controlled drug delivery vehicles that rely on membrane mechanics for cellular entry.

## METHODS

### Cluster size distribution

A necessary condition for the existence of an equilibrium cluster size distribution is the following detailed balance condition [52, 53],

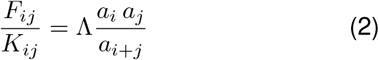

where at this point, Λ and *a*_*k*_’s are arbitrary set of parameters, for which different interpretations exist. One popular approach is the combinatorial interpretation of *k*! *a*_*k*_ as the number of ways that a cluster of size *k* can form from *k* distinguishable particles [52–54]. With **n** = (*n*_1_, *n*_2_, …, *n*_*M*_) being the state vector, with *n*_*k*_ representing number of clusters of size *k*, the equilibrium distribution is given as [52],

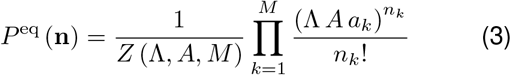

where *A* is the projected surface area, and *Z* (Λ, *A, M*) is the normalizing factor (partition function). The most probable cluster size distribution is thus achieved when,

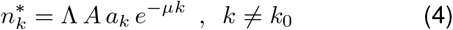

in which the presence of one large cluster of size *k*_0_ is allowed. If this largest cluster incorporates a fraction *g*\* of particles, the mass conservation condition reads as,

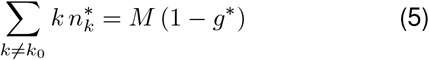

We have assumed *a*_*k*_ ∝ *c*^*k*^, with *c* being a constant. The justification is that the number of ways that a new particle can be added to a 2D cluster of proteins depends on a maximum coordination number. This assumption leads to the following expression for the most probable cluster size distribution,

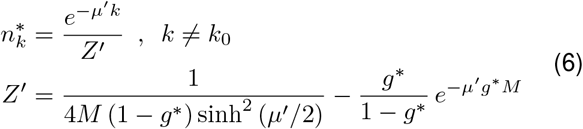

where *μ*′ = *μ* − ln *c* has been substituted. Our assumption for *a*_*k*_ also leads to Λ = *F*_*ij*_*/K*_*ij*_ = *k*_*D*_ (Eqs. (1) and (2)).

### Free energy estimation

We have used the mean of inverse pairwise distances between peripheral proteins as the clustering reaction coordinate,

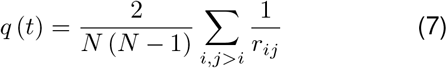

To correctly combine histograms of microstates along *q* from different simulations, we have employed the multiple histogram method [55, 56].

We assume that each simulation, in which a different number of particles have been tagged as proteins, is biased with a Hamiltonian of the form ℋ = ℋ_0_ + *ξ f* (*q*) (Supplementary Information). We justify these assumptions by demonstrating the bias parameter *ξ* to correlate in the two studied cases with (i) the surface concentration (Figs. 3b), or (ii) stiffness of proteins (Fig. 3d).

## SOFTWARE AVAILABILITY

All the in-house developed software used in this study are available from the corresponding authors upon request.

## DATA AVAILABILITY

The data that support the findings of this study are available from the corresponding authors upon reasonable request.

## ACKNOWLEDGEMENTS

The authors wish to thank Thomas Weikl and Patricia Bassereau for comments helpful to the development of this study, as well as Moritz Hoffmann for insights regarding software development.

This research has been funded by Deutsche Forschungsgemeinschaft (DFG) through grants SFB 958/Project A04 “Spatiotemporal model of neuronal signalling and its regulation by presynaptic membrane scaffolds”, SFB 1114/Project C03 “Multiscale modelling and simulation for spatiotemporal master equations”, as well as European Research Commission, ERC CoG 772230 “ScaleCell”.

## AUTHOR CONTRIBUTIONS

MS and FN designed research, MS conducted research and developed software. MS and FN wrote the paper.

## References

[1] Pina, D. G., Johannes, L. & Castanho, M. A. Shiga toxin B-subunit sequential binding to its natural receptor in lipid membranes. Biochim. Biophys. Acta - Biomembr. 1768, 628–636 (2007).

[2] Römer, W. et al. Shiga toxin induces tubular membrane invaginations for its uptake into cells. Nature 450, 670–675 (2007).

[3] Wernick, N. L., Chinnapen, D. J.-F., Cho, J. A. & Lencer, W. I. Cholera toxin: An intracellular journey into the cytosol by way of the endoplasmic reticulum. Toxins (Basel). 2, 310–325 (2010).

[4] Torgersen, M. L., Skretting, G., Van Deurs, B. & Sandvig, K. Internalization of cholera toxin by different endocytic mechanisms. J. Cell Sci. 114, 3737–3747 (2001).

[5] Zimmerberg, J. & Kozlov, M. M. How proteins produce cellular membrane curvature. Nat. Rev. Mol. Cell Biol. 7, 9–19 (2006).

[6] Kozlov, M. M. et al. Mechanisms shaping cell membranes. Curr. Opin. Cell Biol. 29, 53–60 (2014).

[7] Bassereau, P. et al. The 2018 biomembrane curvature and remodeling roadmap. J. Phys. D. Appl. Phys. 51, 343001 (2018).

[8] Watkins, E. B. et al. Shiga Toxin Induces Lipid Compression: A Mechanism for Generating Membrane Curvature. Nano Lett. 19, 7365–7369 (2019).

[9] Weikl, T. R. Membrane-Mediated Cooperativity of Proteins Thomas. Annu. Rev. Phys. Chem 69, 521–539 (2018).

[10] Simunovic, M., Bassereau, P. & Voth, G. A. Organizing membrane-curving proteins: the emerging dynamical picture. Curr. Opin. Struct. Biol. 51, 99–105 (2018).

[11] Windschieg, B. et al. Lipid reorganization induced by Shiga toxin clustering on planar membranes. PLoS One 4, e6238 (2009).

[12] Johannes, L., Pezeshkian, W., Ipsen, J. H. & Shillcock, J. C. Clustering on Membranes: Fluctuations and More. Trends Cell Biol. 28, 405–415 (2018).

[13] Goulian, M., Bruinsma, R. & Pincus, P. Long-range forces in heterogeneous fluid membranes. EPL 23, 125–128 (1993).

[14] Netz, R. R. Inclusions in Fluctuating Membranes: Exact Results. J. Phys. I 7, 833–852 (1997).

[15] Golestanian, R., Goulian, M. & Kardar, M. Fluctuation-induced interactions between rods on membranes and interfaces. Europhys. Lett. 33, 241–245 (1996).

[16] Fournier, J. B. & Dommersnes, P. G. Comment on “Longrange forces in heterogeneous fluid membranes". EPL 39, 681 (1997).

[17] Weikl, T. R., Kozlov, M. M. & Helfrich, W. Interaction of Conical Membrane Inclusions: Effect of Lateral Tension. Phys. Rev. E 57, 10 (1998).

[18] Dommersnes, P. G. & Fournier, J. B. Casimir and meanfield interactions between membrane inclusions subject to external torques. Europhys. Lett. 46, 256–261 (1999).

[19] Schweitzer, Y. & Kozlov, M. M. Membrane-Mediated Interaction between Strongly Anisotropic Protein Scaffolds. PLoS Comput. Biol. 11, e1004054 (2015).

[20] Dommersnes, P. G. & Fournier, J. B. N-body study of anisotropic membrane inclusions: Membrane mediated interactions and ordered aggregation. Eur. Phys. J. B 12, 9–12 (1999).

[21] Reynwar, B. J. et al. Aggregation and vesiculation of membrane proteins by curvature-mediated interactions. Nature 447, 461–464 (2007).

[22] Yoo, J. & Cui, Q. Membrane-mediated protein-protein interactions and connection to elastic models: A coarsegrained simulation analysis of gramicidin a association. Biophys. J. 104, 128–138 (2013).

[23] Weikl, T. R., Hu, J., Xu, G. K. & Lipowsky, R. Binding equilibrium and kinetics of membrane-anchored receptors and ligands in cell adhesion: Insights from computational model systems and theory. Cell Adhes. Migr. 10, 576–589 (2016).

[24] Pezeshkian, W. et al. Mechanism of Shiga Toxin Clustering on Membranes. ACS Nano 11, 314–324 (2017).

[25] Goutaland, Q., Van Wijland, F., Fournier, J. B. & Noguchi, H. Binding of thermalized and active membrane curvature-inducing proteins. Soft Matter 17, 5560–5573 (2021).

[26] Arkhipov, A., Yin, Y. & Schulten, K. Four-scale description of membrane sculpting by BAR domains. Biophys. J. 95, 2806–2821 (2008).

[27] Simunovic, M. et al. Protein-Mediated Transformation of Lipid Vesicles into Tubular Networks. Biophys. J. 105, 711–719 (2013).

[28] Noguchi, H. Membrane tubule formation by banana-shaped proteins with or without transient network structure. Sci. Rep. 6 (2016).

[29] Morlot, S. et al. Membrane shape at the edge of the dynamin helix sets location and duration of the fission reaction. Cell 151, 619–629 (2012).

[30] Schöneberg, J. et al. Lipid-mediated PX-BAR domain recruitment couples local membrane constriction to endocytic vesicle fission. Nat. Commun. (2017).

[31] Schöneberg, J., Ullrich, A. & Noé, F. Simulation tools for particle-based reaction-diffusion dynamics in continuous space. BMC Biophys. 7, 11 (2014).

[32] Hoffmann, M., Fröhner, C. & Noé, F. ReaDDy 2: Fast and flexible software framework for interacting-particle reaction dynamics. PLoS Comput. Biol. 15, e1006830 (2019).

[33] Fröhner, C. & Noé, F. Reversible Interacting-Particle Reaction Dynamics. J. Phys. Chem. B 122, 11240–11250 (2018).

[34] Dibak, M., del Razo, M. J., De Sancho, D., Schütte, C. & Noé, F. MSM/RD: Coupling Markov state models of molecular kinetics with reaction-diffusion simulations. J. Chem. Phys. 148, 214107 (2018).

[35] del Razo, M. J., Dibak, M., Schütte, C. & Noé, F. Multiscale molecular kinetics by coupling Markov state models and reaction-diffusion dynamics. 2103.06889 (2021).

[36] Sadeghi, M. & Noé, F. Large-scale simulation of biomem-branes incorporating realistic kinetics into coarse-grained models. Nat. Commun. 11, 2951 (2020).

[37] Frenkel, D. & Smit, B. Understanding Molecular Simulation: From Algorithms to Applications (Academic Press, 2002), 2nd edn.

[38] Levashov, V. A., Morris, J. R. & Egami, T. Viscosity, shear waves, and atomic-level stress-stress correlations. Phys. Rev. Lett. 106, 2–5 (2011).

[39] Sadeghi, M., Weikl, T. R. & Noé, F. Particle-based membrane model for mesoscopic simulation of cellular dynamics. J. Chem. Phys. 148, 044901 (2018).

[40] Pezeshkian, W. et al. Membrane invagination induced by Shiga toxin B-subunit: from molecular structure to tube formation. Soft Matter 12, 5164–5171 (2016).

[41] Tian, A. & Baumgart, T. Sorting of lipids and proteins in membrane curvature gradients. Biophys. J. 96, 2676–2688 (2009).

[42] Höfling, F. & Franosch, T. Anomalous transport in the crowded world of biological cells. Reports Prog. Phys. 76 (2013).

[43] Day, C. A. & Kenworthy, A. K. Mechanisms underlying the confined diffusion of cholera toxin B-subunit in intact cell membranes. PLoS One 7, e34923 (2012).

[44] Knight, J. D., Lerner, M. G., Marcano-Velázquez, J. G., Pastor, R. W. & Falke, J. J. Single molecule diffusion of membrane-bound proteins: Window into lipid contacts and bilayer dynamics. Biophys. J. 99, 2879–2887 (2010).

[45] Waugh, R. E. Surface viscosity measurements from large bilayer vesicle tether formation. II. Experiments. Biophys. J. 38, 29–37 (1982).

[46] Dimova, R., Dietrich, C., Hadjiisky, a., Danov, K. & Pouligny, B. Falling ball viscosimetry of giant vesicle membranes: Finite-size effects. Eur. Phys. J. B 12, 589–598 (1999).

[47] Camley, B. A., Esposito, C., Baumgart, T. & Brown, F. L. H. Lipid Bilayer Domain Fluctuations as a Probe of Membrane Viscosity. Biophys. J. 99, L44–L46 (2010).

[48] Faizi, H. A., Dimova, R. & Vlahovska, P. M. Viscosity of fluid membranes measured from vesicle deformation. bioRxiv 433848 (2021).

[49] Zgorski, A., Pastor, R. W. & Lyman, E. Surface Shear Viscosity and Interleaflet Friction from Nonequilibrium Simulations of Lipid Bilayers. J. Chem. Theory Comput. 15, 6471–6481 (2019).

[50] Shkulipa, S. a., Den Otter, W. K. & Briels, W. J. Surface viscosity, diffusion, and intermonolayer friction: simulating sheared amphiphilic bilayers. Biophys. J. 89, 823–829 (2005).

[51] Ford, I. J. Statistical mechanics of nucleation: A review. Proc. Inst. Mech. Eng. Part C J. Mech. Eng. Sci. 218, 883–899 (2004).

[52] Hendriks, E. M. Cluster size distributions in equilibrium. Zeitschrift fü r Phys. B Condens. Matter 57, 307–314 (1984).

[53] van Dongen, P. G. & Ernst, M. H. Kinetics of reversible polymerization. J. Stat. Phys. 37, 301–324 (1984).

[54] Spouge, J. L. A branching-process solution of the polydisperse coagulation equation. Adv. Appl. Probab. 16, 56–69 (1984).

[55] Ferrenberg, A. M. & Swendsen, R. H. Optimized Monte Carlo data analysis. Phys. Rev. Lett. 63, 1195–1198 (1989).

[56] Kumar, S., Rosenberg, J. M., Bouzida, D., Swendsen, R. H. & Kollman, P. A. The weighted histogram analysis method for free energy calculations on biomolecules. I. The method. J. Comput. Chem. 13, 1011–1021 (1992).

