## Supplementary Information for "Thermodynamics and kinetics of aggregation of flexible peripheral membrane proteins"

### A. Elastic model of membrane and peripheral proteins

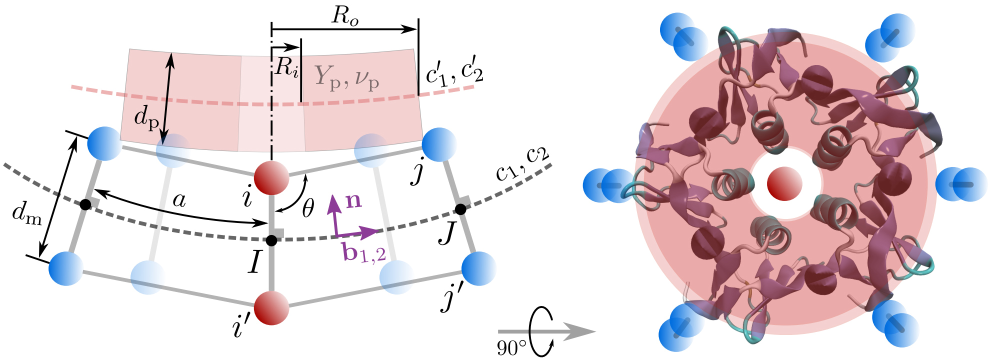


Figure 1: Schematic of the membrane model where a peripheral protein is binding. The discrete geometry of the particle-based model reflects the curvature induced in the membrane. The governing force field is parameterized to reproduce the combined elasticity of the membrane and the protein.

For a homogeneous membrane the elastic energy is very well described by the Helfrich model [1–4],

$$f_{\mathrm{H}}(\mathbf{x}) = 2\kappa \left(H(\mathbf{x}) - H_0(\mathbf{x})\right)^2 + \bar{\kappa} G(\mathbf{x}) \tag{1}$$

in which $H(\mathbf{x}) = \frac{1}{2}(c_1 + c_2)$ and $G(\mathbf{x}) = c_1 c_2$ are the mean and the Gaussian curvatures of the membrane at position $\mathbf{x}$, with $c_1$ and $c_2$ being the principal curvatures (Fig. 1). $\kappa$ and $\bar{\kappa}$ are the corresponding bending rigidity and Gaussian curvature moduli, and $H_0(\mathbf{x})$ is the spontaneous curvature, arising from asymmetries between the two leaflets and the shapes of individual lipid molecules.

To build an elastic model of the peripheral protein, we assume an intrinsic curvature, $c_0$, for the protein, such that the bound protein is unstrained when the underlying membrane patch acquires the mean inward curvature of $c_0$. But the flexibility of the protein as well as the protein-membrane coupling means that in mechanical equilibrium, the protein has non-zero principal curvatures $c_1'$ and $c_2'$. The curvatures of the protein and the membrane are complementary in the sense that $c_1' + c_1 = c_0$ and $c_2' + c_2 = c_0$ (Fig. 1). We employ the Kirchhof-Love small-deformation plate theory as a continuum model of the protein, resulting in the protein strain energy per unit area,

$$f_{\mathrm{p}}(\mathbf{x})=\frac{Y_{\mathrm{p}}d_{\mathrm{p}}^{3}}{6\left(1-\nu_{\mathrm{p}}^{2}\right)}\left[\left(H(\mathbf{x})-c_{0}\right)^{2}-\frac{1-\nu_{\mathrm{p}}}{2}\left(c_{1}(\mathbf{x})-c_{0}\right)\left(c_{2}(\mathbf{x})-c_{0}\right)\right] \tag{2}$$

in which $Y_{\mathrm{p}}$ is the effective Young modulus of the protein, $\nu_{\mathrm{p}}$ is its Poisson's ratio and $d_{\mathrm{p}}$ is its thickness (Fig. 1). Having the energy densities of both the protein and the membrane, the total energy density for which the particle-based model will be parametrized is,

$$f_{\mathrm{total}}(\mathbf{x})=f_{\mathrm{H}}(\mathbf{x})+\mathbb{1}_{\mathrm{p}}(\mathbf{x})\,f_{\mathrm{p}}(\mathbf{x})\frac{A_{\mathrm{p}}}{A_{\mathrm{m}}} \tag{3}$$

where $\frac{A_{\mathrm{p}}}{A_{\mathrm{m}}}$ denotes the surface area ratio between the peripheral protein and the membrane in the binding region, and $\mathbb{1}_{\mathrm{p}}(\mathbf{x})$ is an indicator function equal to one where a peripheral protein is bound to the membrane, and zero otherwise.

### B. Particle-based model of membrane and peripheral proteins

The membrane model used for the simulations presented here, is the same as developed in [5] and used in [6, 7]. The bilayer is modeled as formed by particle-dimers in a close-packed arrangement, with a lattice parameter of a few nanometers, while the two leaflets are resolved (Fig. 1)). Bonded interactions in the model are described via the following potentials [5],

$$U_{\mathrm{s}}(r_{ij})=D_{e}\left[1-\exp\left(-\alpha\left(r_{ij}-r_{\mathrm{eq}}\right)\right)\right]^{2} \tag{4a}$$

$$U_{\mathrm{a}}(\theta_{i'ij})=K_{a}\left(\theta_{i'ij}-\theta_{\mathrm{eq}}\right)^{2} \tag{4b}$$

$$U_{\mathrm{d}}(d_{ii'})=K_{d}\left(d_{ii'}-d_{\mathrm{eq}}\right)^{2} \tag{4c}$$

Particles belonging to each leaflet are connected to their nearest-neighbour counterparts via Morse-type bonds (Eq. (4a)). Also, harmonic angle-bending potentials given by Eq. (4b) act against the out-of-plane rotations of these bonds. Finally, particles in a dimer are connected via harmonic bonds of the form described by Eq. (4c), which keeps the two leaflets together (Fig. 1)).

The force field masking in the presence of the protein is done such that the underlying membrane model be least affected. For potentials in Eqs. (4), we modify $r_{\mathrm{eq}}$ and $\theta_{\mathrm{eq}}$ for bonds in top and bottom leaflets. $D_{\mathrm{e}}$ and $\alpha$ are left unchanged, leading to $\alpha^{2}/D_{\mathrm{e}}$, the stiffness of the Morse bond at $r_{\mathrm{eq}}$, to remain constant. The rest of the effects are implemented via the auxiliary angle-bending potential:

$$U'_{\mathrm{a}}\left(\theta_{i'ij}\right) = K'_{a}\left(\theta_{i'ij} - \theta_{\mathrm{eq}}\right)^{2} \tag{5}$$

which reinforces the angle-bending potentials on the top leaflet for the masked particles.

### C. Differential geometry of the particle-based model

Consider a locally tangent orthonormal basis constructed at point $I$ on the mid-surface of the membrane (Fig. 1)). Position vectors in this basis are given as $\mathbf{r} = x\,\mathbf{b}_1^I + y\,\mathbf{b}_2^I + z\,\mathbf{n}^I$, in which, $\mathbf{n}^I = \mathbf{b}_1^I \times \mathbf{b}_2^I$ is the surface normal at $I$. The mid-surface of the membrane can be approximated to the second order through an osculating paraboloid. Assuming the two principal curvatures of the mid-surface to be $c_1$ and $c_2$, and the base vector $\mathbf{b}_1^I$ making an angle $\phi$ with the principal direction corresponding to $c_1$, the osculating paraboloid is defined via the following relation,

$$\begin{aligned} 2z &= Ax^2 + 2Bxy + Cy^2 \\ A &= H + \frac{c_1 - c_2}{2}\cos 2\phi \\ B &= \frac{c_1 - c_2}{2}\sin 2\phi \\ C &= H - \frac{c_1 - c_2}{2}\cos 2\phi \end{aligned} \tag{6}$$

It can be readily verified that the mean and Gaussian curvatures are given as $H = \frac{1}{2}\left(A + C\right)$ and $G = AC - B^2$. We assume the projection of the bond between particles $i$ and $j$ to be parallel to $\mathbf{b}_1$ (which can always be achieved by a change in the angle $\phi$). To find the point $J$, the projection of the position of particle $j$ on the mid-surface, we assume the mid-surface to be incompressible. Thus, along the $y = 0$ section of the osculating paraboloid, we can find the projection of point $j$, given by the coordinates $(x_J, 0, z_J)$, assuming it was a distance $a$ apart from $I$ in the undeformed configuration,

$$a = \int_0^{x_J} \sqrt{1 + \left(\frac{\partial z}{\partial x}\right)^2_{y=0}}\, dx = x_J + \frac{A^2}{6}x_J^3 + \mathcal{O}\left(x_J^5\right) \tag{7}$$

discarding $x_J^5$ and higher-order terms, a closed-form solution is possible. Based on the positions of points $I$ and $J$, positions of particles $i$ and $j$ are given as,

$$\begin{aligned}\mathbf{r}_i &= \mathbf{r}_I + \frac{d_\mathrm{m}}{2}\mathbf{n}^I \\ \mathbf{r}_j &= \mathbf{r}_J + \frac{d_\mathrm{m}}{2}\mathbf{n}^J \\ &= \mathbf{r}_I + x_J\mathbf{b}_1^I + \frac{Ax_J^2}{2}\mathbf{n}^I + \frac{d_\mathrm{m}}{2}\mathbf{n}^J\end{aligned} \tag{8}$$

where $d$ is the thickness of the membrane and

$$\mathbf{n}_J = \frac{\left(-Ax_J\,\mathbf{b}_1^I - Bx_J\,\mathbf{b}_2^I + \mathbf{n}^I\right)}{\sqrt{1 + A^2x_J^2 + B^2x_J^2}} \tag{9}$$

Having the position of the particle $j$ in the neighborhood of particle $i$, we can relate the bond-stretching potential between the two (Eq. (4a)) to the curvature of the mid-surface. Similarly, for the out-of-plane angle-bending potential (Eq. (4b)) the triplet of particles $i'\,i\,j$ is considered, with $\mathbf{r}_{i'} = \mathbf{r}_I - \frac{d_\mathrm{m}}{2}\mathbf{n}_I$ (Fig. 1).

### D. Force field parametrization

The force field attributed to bonded interactions results in an effective energy density, $f_\mathrm{eff}$. We have estimated this energy density as the ratio of all the potential energies attributed to one particle to the area per particle in the model. We have also averaged this ratio over $\phi$ in Eq. (6) to eliminate the effect due to the angle between bonds and the principal directions of the membrane [5]. The force field parameters are obtained via minimizing the loss function,

$$\mathcal{L} = \mathop{\mathbb{E}}_{c_1,c_2}\left[\left(f_\mathrm{eff} - f_\mathrm{tot}\right)^2\right] + \sum_i \lambda_i \mathop{\mathbb{E}}_{c_1,c_2}\left[\left(X_{\mathrm{eff},i} - X_0\right)^2\right] \tag{10}$$

where $X_{\mathrm{eff},i}$'s are other properties for which an effective value can be obtained in the space of inspected membrane geometries.

We chose the Broyden-Fletcher-Goldfarb-Shanno algorithm (BFGS) for parameter-space optimization [8], using the implementation included in SciPy optimization library [9]. The input physical properties of the membrane and flexible proteins are listed in Tab. I. The values of the force field parameters, obtained or chosen for simulations, are summarized in Tab. II.

Table I: Properties of the membrane and peripheral proteins used for the force field parametrization [10–17].

| Membrane | | | | | |
|---|---|---|---|---|---|
| $d_{\mathrm{m}}$ [nm] | $\kappa$ [$kT$] | $\bar{\kappa}$ [$kT$] | $K_{\mathrm{area}}$ $\left[\mathrm{N\,m^{-1}}\right]$ | | |
| 4.0 | 18.73 | $-14.98$ | 0.270 | | |
| **Peripheral protein** | | | | | |
| $d_{\mathrm{p}}$ [nm] | $R_i$ [nm] | $R_o$ [nm] | $c_0$ $\left[\mathrm{nm^{-1}}\right]$ | $Y_{\mathrm{p}}$ [MPa] | $\nu_{\mathrm{p}}$ |
| 2.92 | 0.49 | 3.21 | 0.08 | 50, 100, 200 | 0.25 |

### E. Simulations and analysis

We have used over-damped Langevin dynamics with anisotropic diffusion tensors to describe the motion of membrane and protein particles [6, 7]. We have considered hydrodynamic coupling to an aqueous solvent with the viscosity of $\eta = 0.890\,\mathrm{mPa\,s}$, and have included hydrodynamic interactions between nearest-neighbor particles (the "Hydro NN" model described in ref. [6]). All simulations have been carried out at the biological temperature of $T = 310\,\mathrm{K}$. We have coupled the in-plane degrees of freedom to the Langevin piston barostat to simulate tensionless membranes [18].

The in-plane fluidity of the membrane is modeled using bond-flipping Monte Carlo moves [5]. One move constitutes switching a bond shared between two adjacent bonded triangles to the crossing diagonal. We evaluate the acceptance probability of a proposed bond-flip, as well as force field masking due to protein presence, via the Metropolis-Hastings algorithm [19]. In order to guarantee that the masking of the force field does not disturb bond-flipping, as described in Sec. C, we have separated the masking of the force field into two distinct contributions, one that preserves the stiffness of bonds, and an additional stiffening potential. Bond-flipping is decided based on the former set of interactions, and results in the force field parameters of flipped bonds to reset to the original unmasked values. We have monitored the potential energy of bonded interactions throughout the simulations to ensure that this process does not result in any unwanted energy fluctuations.

To measure the effective local curvature around each protein-supporting particle (main

Table II: Force field parameters for the interaction potentials given in Eqs. (4) and (5).

| Unmasked (membrane) | | | |
|---|---|---|---|
| $r_{\mathrm{eq}}$ [nm] | $D_{\mathrm{e}}$ $\left[\mathrm{kJ\,mol^{-1}}\right]$ | $\alpha$ $\left[\mathrm{nm^{-1}}\right]$ | |
| 6.5 | 9.03 | 0.18 | |
| $\theta_{\mathrm{eq}}$ [rad] | $K_{\mathrm{a}}$ $\left[\mathrm{kJ\,mol^{-1}}\right]$ | | |
| $\pi/2$ | 19.35 | | |
| $d_{\mathrm{eq}}$ [nm] | $K_{\mathrm{d}}$ $\left[\mathrm{kJ\,mol^{-1}nm^{-2}}\right]$ | | |
| 4.0 | 6.19 | | |
| Masked (membrane + protein) | | | |
| $Y_{\mathrm{p}} = 50\,\mathrm{MPa}$ | | | |
| | $r'_{\mathrm{eq}}$ [nm] | $\theta'_{\mathrm{eq}}$ [rad] | $K'_{\mathrm{a}}$ $\left[\mathrm{kJ\,mol^{-1}}\right]$ |
| top leaflet: | 5.96 | 1.70 | 85.38 |
| bottom leaflet: | 7.00 | 1.44 | - |
| $Y_{\mathrm{p}} = 100\,\mathrm{MPa}$ | | | |
| | $r'_{\mathrm{eq}}$ [nm] | $\theta'_{\mathrm{eq}}$ [rad] | $K'_{\mathrm{a}}$ $\left[\mathrm{kJ\,mol^{-1}}\right]$ |
| top leaflet: | 5.88 | 1.72 | 160.37 |
| bottom leaflet: | 7.07 | 1.42 | - |
| $Y_{\mathrm{p}} = 200\,\mathrm{MPa}$ | | | |
| | $r'_{\mathrm{eq}}$ [nm] | $\theta'_{\mathrm{eq}}$ [rad] | $K'_{\mathrm{a}}$ $\left[\mathrm{kJ\,mol^{-1}}\right]$ |
| top leaflet: | 5.67 | 1.78 | 232.20 |
| bottom leaflet: | 7.24 | 1.37 | - |

Fig. 1b), we have fitted a bivariate quadratic function to the position of each particle and its nearest neighbors. These spontaneous values of curvatures have been sampled for a random selection of particles for each frame of the trajectory.

To obtain cluster size information from trajectories, we have used the Density-Based Spatial Clustering of Applications with Noise (DBSCAN) algorithm [20]. This method can discover point clusters based on a pairwise distance metric and a tolerance measure for particles belonging to a cluster. We have set the tolerance value to be 1.5 times the lattice parameter of the model.

For free energy estimation, with the suggested Hamiltonian of the form $\mathcal{H} = \mathcal{H}_0 + \xi f(q)$, we have estimated the function $f$ with a polynomial. We obtained the optimal polynomial as well as the values of the bias $\xi$ via minimizing the mean absolute error in the estimation of $\mathcal{H}$. We used the SciPy implementation of the Nelder-Mead method for the optimization [9, 21].

### F. Software

We have performed simulations based on the particle-based membrane model [5–7] using our in-house specific-purpose C++ software. We have used multithreading for parallel computation with enhanced performance. Python libraries Numpy, SciPy and Matplotlib are used for analysis and presentation of the results [9, 22, 23]. The software package Visual Molecular Dynamics (VMD) is used for some visualizations [24].


[1] Helfrich, W. Elastic Properties of Lipid Bilayers: Theory and Possible Experiments. *Zeitschrift fur Naturforsch. - Sect. C J. Biosci.* **28**, 693–703 (1973).

[2] Canham, P. B. The minimum energy of bending as a possible explanation of the biconcave shape of the human red blood cell. *J. Theor. Biol.* **26**, 61–81 (1970).

[3] Evans, E. A. Bending resistance and chemically induced moments in membrane bilayers. *Biophys. J.* **14**, 923–931 (1974).

[4] Campelo, F., Arnarez, C., Marrink, S. J. & Kozlov, M. M. Helfrich model of membrane bending: From Gibbs theory of liquid interfaces to membranes as thick anisotropic elastic layers. *Adv. Colloid Interface Sci.* **208**, 25–33 (2014).

[5] Sadeghi, M., Weikl, T. R. & Noé, F. Particle-based membrane model for mesoscopic simulation of cellular dynamics. *J. Chem. Phys.* **148**, 044901 (2018).

[6] Sadeghi, M. & Noé, F. Large-scale simulation of biomembranes incorporating realistic kinetics into coarse-grained models. *Nat. Commun.* **11**, 2951 (2020).

[7] Sadeghi, M. & Noé, F. Hydrodynamic coupling for particle-based solvent-free membrane models. *arXiv* 1909.02722 (2020).

[8] Liu, D. C. & Nocedal, J. On the limited memory BFGS method for large scale optimization. *Math. Program.* **45**, 503–528 (1989).

[9] Virtanen, P. *et al.* SciPy 1.0: fundamental algorithms for scientific computing in Python. *Nat. Methods* **17**, 261–272 (2020).

[10] Marsh, D. Elastic curvature constants of lipid monolayers and bilayers. *Chem. Phys. Lipids* **144**, 146–159 (2006).

[11] Hu, M., Briguglio, J. J. & Deserno, M. Determining the Gaussian curvature modulus of lipid membranes in simulations. *Biophys. J.* **102**, 1403–1410 (2012).

[12] Nagle, J. F. Introductory Lecture: Basic quantities in model biomembranes. *Faraday Discuss.* **161**, 11–29 (2013).

[13] Dimova, R. Recent developments in the field of bending rigidity measurements on membranes. *Adv. Colloid Interface Sci.* **208**, 225–234 (2014).

[14] Janosi, L. & Gorfe, A. A. Simulating POPC and POPC/POPG bilayers: Conserved packing and altered surface reactivity. *J. Chem. Theory Comput.* **6**, 3267–3273 (2010).

[15] Klauda, J. B. *et al.* Update of the CHARMM All-Atom Additive Force Field for Lipids: Validation on Six Lipid Types. *J. Phys. Chem. B* **114**, 7830–7843 (2010).

[16] Raghunathan, M. *et al.* Structure and elasticity of lipid membranes with genistein and daidzein bioflavinoids using X-ray scattering and MD simulations. *J. Phys. Chem. B* **116**, 3918–3927 (2012).

[17] Braun, A. R., Sachs, J. N. & Nagle, J. F. Comparing simulations of lipid bilayers to scattering data: The GROMOS 43A1-S3 force field. *J. Phys. Chem. B* **117**, 5065–5072 (2013).

[18] Feller, S. E., Zhang, Y., Pastor, R. W. & Brooks, B. R. Constant pressure molecular dynamics simulation: The Langevin piston method. *J. Chem. Phys.* **103**, 4613–4621 (1995).

[19] Frenkel, D. & Smit, B. *Understanding Molecular Simulation: From Algorithms to Applications* (Academic Press, 2002), 2nd edn.

[20] Ester, M., Kriegel, H.-P., Sander, J. & Xu, X. A Density-Based Algorithm for Discovering Clusters in Large Spatial Databases with Noise. In *Proc. 2nd Int. Conf. Knowl. Discov. Data Min.*, 226–231 (1996).

[21] Nelder, J. A. & Mead, R. A Simplex Method for Function Minimization. *Comput. J.* **7**, 308–313 (1965).

[22] Harris, C. R. *et al.* Array programming with NumPy. *Nature* **585**, 357–362 (2020).

[23] Hunter, J. D. Matplotlib: A 2D graphics environment. *Comput. Sci. Eng.* **9**, 90–95 (2007).

[24] Humphrey, W., Dalke, A. & Schulten, K. VMD - Visual Molecular Dynamics. *J. Molec. Graph.* **14**, 33–38 (1996).
